# Parietal Mechanisms for Transsaccadic Spatial Frequency Perception: An fMRI Study

**DOI:** 10.1101/2020.07.14.203190

**Authors:** B. R. Baltaretu, B. T. Dunkley, W. Dale Stevens, J. D. Crawford

**Affiliations:** Centre for Vision Research and Vision: Science to Applications (VISTA) program, York University, Toronto, Ontario M3J 1P3, Canada; Department of Biology, York University, Toronto, Ontario M3J 1P3, Canada; Department of Diagnostic Imaging, Hospital for Sick Children, Toronto, Ontario M5G 1X8, Canada; Neurosciences & Mental Health, Hospital for Sick Children Research Institute, Toronto, Ontario M5G 1X8, Canada; Department of Medical Imaging, University of Toronto, Toronto, Ontario M5S, Canada; Department of Psychology and Neuroscience Graduate Diploma Program, York University, Toronto, Ontario M3J 1P3, Canada; School of Kinesiology and Health Sciences, York University, Toronto, Ontario M3J 1P3, Canada

## Abstract

Posterior parietal cortex (PPC), specifically right supramarginal gyrus, is involved in transsaccadic memory of object orientation for both perception and action. Here, we investigated whether PPC is involved in transsaccadic memory of other features, namely spatial frequency. We employed a functional magnetic resonance imaging paradigm where participants briefly viewed a grating stimulus with a specific spatial frequency that later reappeared with the same or different frequency, after a saccade or continuous fixation. Post-saccadic frequency modulation activated a region in the right hemisphere spanning medial PPC (ventral precuneus) and posterior cingulate cortex. Importantly, the site of peak precuneus activation showed saccade-specific feature modulation (compared to fixation) and task-specific saccade modulation (compared to a saccade localizer task). Psychophysiological interaction analysis revealed functional connectivity between this precuneus site and the precentral gyrus (M1), lingual gyrus (V1/V2), and medial occipitotemporal sulcus. This differed from the transsaccadic *orientation* network, perhaps because spatial frequency signaled changes in object *identity*. Overall, this experiment supports a general role for PPC in transsaccadic vision, but suggests that different networks are employed for specific features.

## Introduction

The visual system tracks both low-level (e.g., orientation, spatial frequency) and high-level (e.g., objects, faces) components of our visual surroundings through space and time [1], despite the interruption of several saccades (rapid eye movements) per second [2,3]. To do this, visual features must be encoded, retained, updated, and integrated across saccades [4,5], through a process called transsaccadic perception [2,6,7,8]. As argued elsewhere [9–11], transsaccadic perception likely incorporates mechanisms for both visual working memory [12,13] and spatial updating [14,15]. However, the specific *neural* mechanisms for transsaccadic feature perception are not well understood.

When saccades occur, they cause both object locations and their associated features to shift relative to eye position. It is well established that human posterior parietal cortex (PPC; specifically, the mid-posterior parietal sulcus), is involved in transsaccadic spatial updating, i.e., the updating of object *location* relative to each new eye position [16–19]. Recently, we found that inferior PPC (specifically, right supramarginal gyrus; SMG) is also modulated by transsaccadic comparisons of object orientation [20]. This area is located immediately lateral to the parietal eye field / lateral intraparietal sulcus, so might be an evolutionary expansion of the monkey intraparietal eye fields, which show modest spatial updating of object information across saccades [21]. Consistent with these findings, transcranial magnetic stimulation (TMS) of PPC (just posterior to SMG) disrupted transsaccadic memory of multiple object orientations [22,23]. SMG activity is also modulated during transsaccadic updating of object orientation for grasp, along with other parietal sensorimotor areas [24]. These findings implicate PPC in the transsaccadic updating of *both* object location and orientation.

It is not known if these neural mechanisms generalize to other stimulus features. One might expect PPC to be involved in other aspects of transsaccadic feature memory and integration, because of its general role in spatial updating [17,25,26]; however, the specific mechanisms might differ. SMG seems to play a specialized role for high-level object orientation in various spatial tasks [27,28]. Thus, while SMG might play a general role in transsaccadic updating of all visual features, it is equally possible that the brain engages different cortical networks for transsaccadic processing of different features, as it does during prolonged visual fixations [29,30].

To address this question, we used an event-related fMRI paradigm, similar to Dunkley et al. [20], where participants briefly viewed a 2D spatial frequency grating stimulus, either while continually fixating the eyes, or while making a saccade to the opposite side, and then judged whether a re-presented grating was the same or different. But here, we modulated spatial frequency, rather than orientation (Fig. 1a,b). As in Dunkley et al. [20], we first localized clusters that were activated by changes in stimulus frequency following saccades; then, as in Baltaretu et al. [24] we determined if the site(s) of peak PPC activation passed two additional criteria: their feature modulations must be saccade-specific (Fig. 2, prediction 1), and 2) they must show task-specific saccade modulations relative to a simple saccade motor task (Fig. 2, prediction 2). Finally, we analyzed the functional connectivity of this area. Our findings confirm the involvement of PPC in processing transsaccadic feature interactions, but demonstrate that different PPC areas/networks are involved in transsaccadic processing of spatial frequency versus orientation.

**Figure 1.**
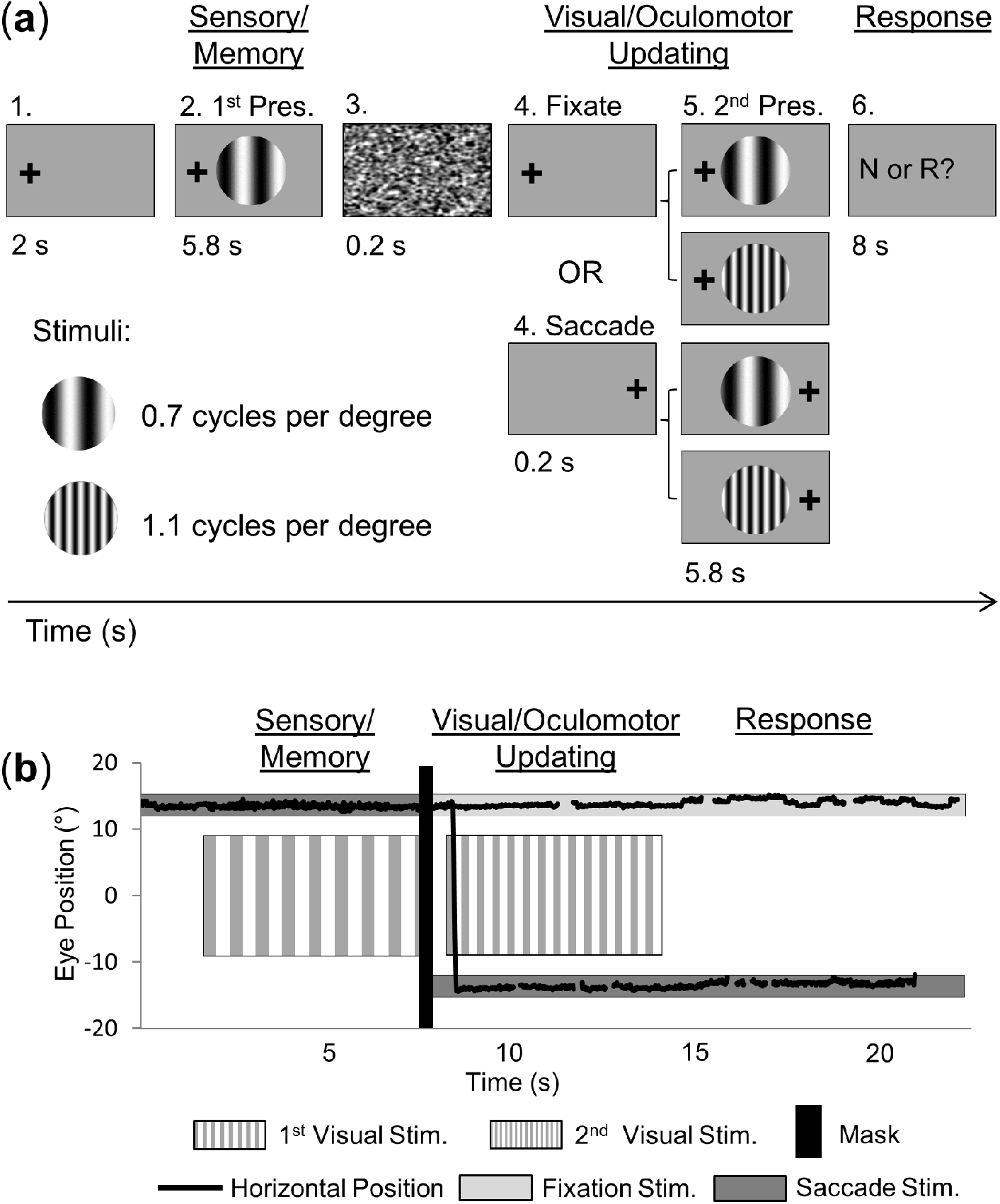
Experimental paradigm and eye movement traces. **A.** An example trial is shown (0.7 cycles per degree, cpd, left fixation) with the four possible conditions: Fixation Same, Fixation Different, Saccade Same, and Saccade Different. Each 22 s trial had three major phases: 1) *Sensory/Motor*, for the first presentation of the stimulus at one of the two possible spatial frequencies (0.7 or 1.1 cpd) while gaze could be to the left or right; 2) *Visual/Oculomotor Updating*, for the second presentation of the stimulus at the same spatial frequency (e.g., 0.7 cpd for first and second stimulus presentations; *Same* condition) or different (e.g., 0.7 cpd for first presentation and 1.1 cpd for second stimulus presentation, or vice versa; *Different* condition) while participants maintained fixation on the same cross *(Fixate* condition) or made a directed saccade *(Saccade* condition); and 3) *Response*, for the button press response period where an indication of whether the spatial frequency across the two stimulus presentations was the same (‘R’) or different (‘N’). **B.** An example eye position trace (°) for an example fixation and saccade trial. In this figure, each of the two trials started with initial fixation on the right (13.5° from center), then diverged after the initial presentation of the stimulus (1^st^ visual stimulus) and mask (black vertical bar), whereby gaze remained fixed for the fixation trial or moved to the other fixation cross position (−13.5° from center) for the saccade trial, where it remained after the second stimulus presentation (2^nd^ visual stimulus), and button press period.

**Figure 2.**
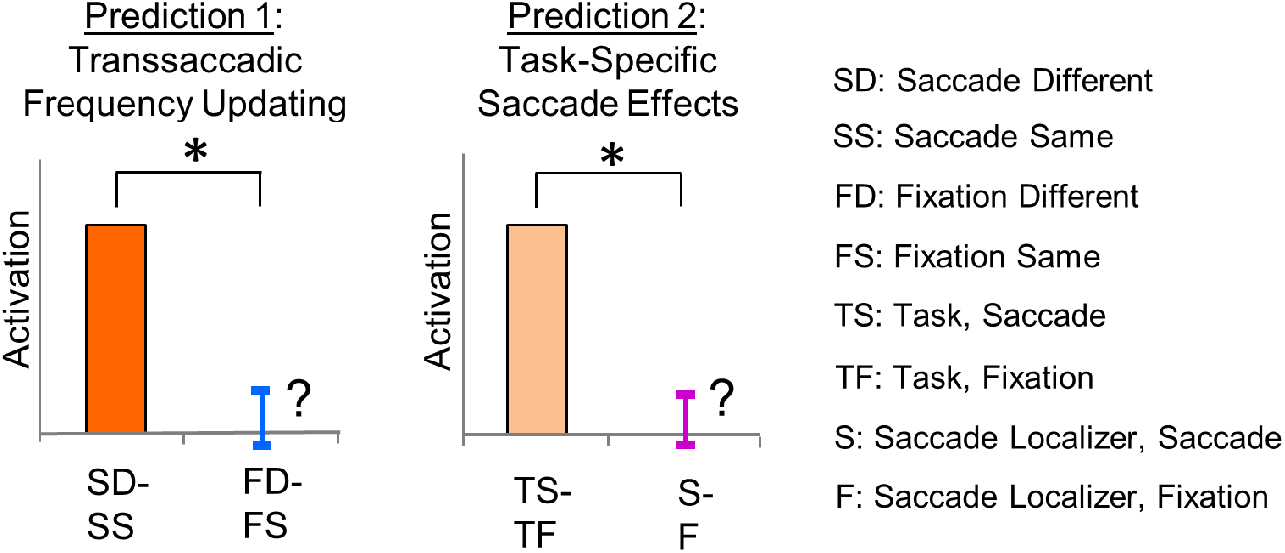
Predictions used to test transsaccadic feature- and task-specificity. Prediction 1: For an area that shows saccade-related feature-specific modulations, there should be a larger response 1) for a change in spatial frequency than a repetition, and 2) when a saccade is produced than for fixation (i.e., (Saccade Different – Saccade Same) > (Fixation Different – Fixation Same); SD, SS, FD, FS, respectively). Prediction 2: The saccade-related effects in this region should also show task-specificity – i.e., there should be larger modulations in the Saccade condition than the Fixation condition for the experimental task compared with the independent saccade localizer (i.e., (Task, Saccade – Task, Fixation) > (Saccade Localizer, Saccade – Saccade Localizer, Fixation); TS, TF, S, F, respectively).

## Results

Overall, our task (Fig. 1) produced widespread and progressive activation in brain areas related to vision, visual memory and eye movements during the initial *Sensory/Memory Phase* (Fig. S1) and *Visual/Oculomotor Updating Phase* (Fig. S2). Here, we focus on the specific analysis pipeline that we used to identify and test if any parietal sites showed saccade-and task-specific feature modulations for spatial frequency (Fig. 2). For this analysis, we only analyzed the *Visual/Oculomotor Updating* phase of our task, i.e. the period after the first stimulus presentation, starting at the time when a saccade occurred and a second stimulus (either Same or Different) appeared (Fig. 1a). An *a priori* power analysis suggested that 14 participants were required for the voxelwise contrasts used in this pipeline (see Methods: Power analysis). To obtain this level, we continued testing participants (21 in total), until 15 of these passed our behavioural inclusion criteria for fMRI analysis (See Methods: Behavioural data and exclusion criteria).

### Selection of parietal site for hypothesis testing

Our overall aim was to identify parietal site(s) that showed post-saccadic feature modulations and then use these sites to test the predictions shown in Figure 2. As a first step in this analysis, we performed a whole-brain voxelwise contrast (*Different* Spatial Frequency > *Same* Spatial Frequency) on fMRI data derived from the *Visual/Oculomotor Updating* phase of our saccade trials. The resulting group data are shown in Fig. 3a, overlaid on a representative anatomic scan in ‘inflated brain’ coordinates (note that while visually convenient, this convention results in small spatial distortions). This contrast revealed two regions: First, a continuous region of change-dependent activation in the right hemisphere spanning medial PPC (ventral precuneus and the posterior cingulate gyrus, PCu/pCG). The other region (right medial frontal gyrus, MFG), showed the opposite pattern, i.e., change suppression, but was not further analyzed. Notably, unlike similar contrasts in studies of transsaccadic *orientation* processing [20], SMG was not significantly modulated.

**Figure 3.**
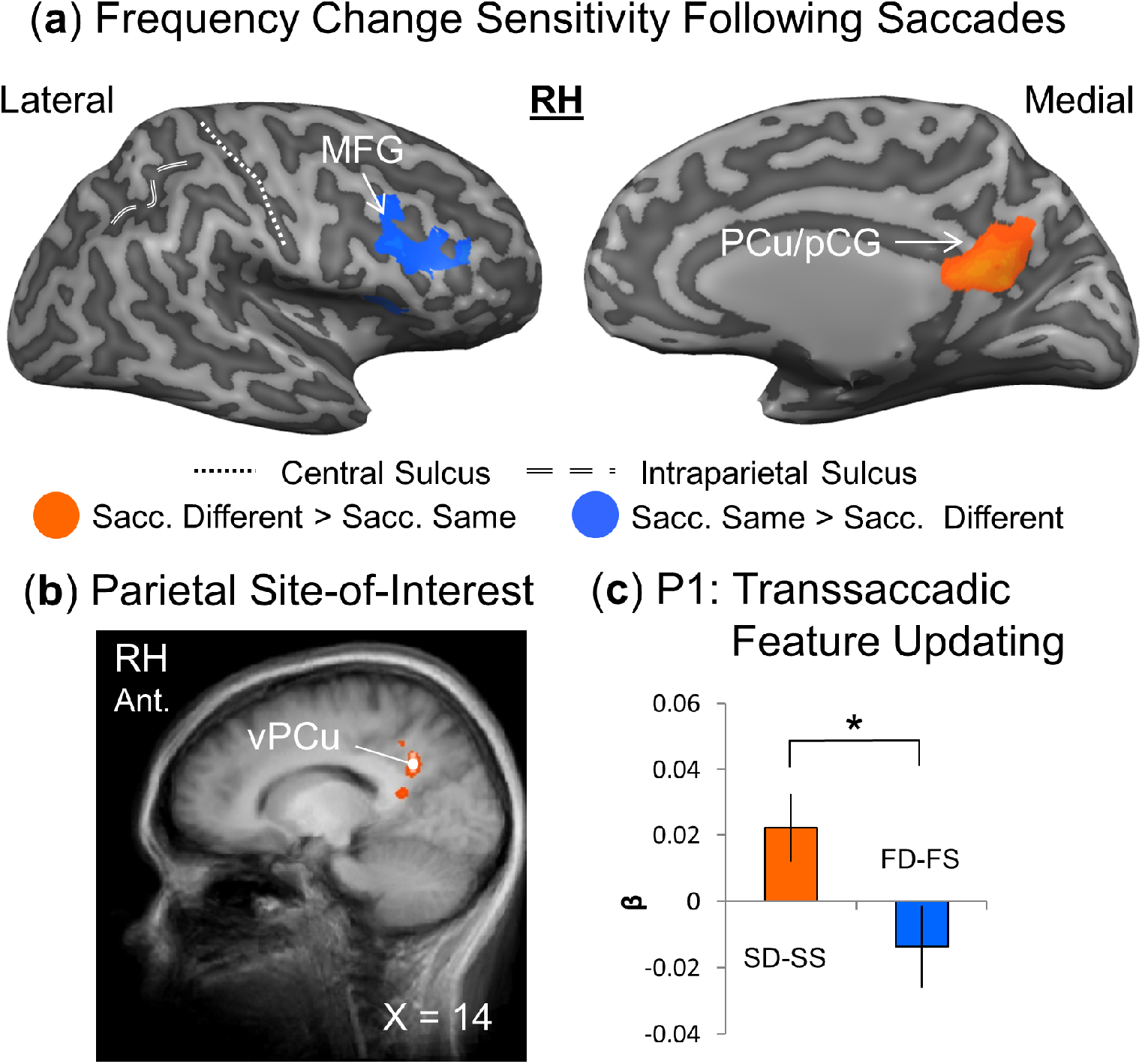
Testing Prediction 1: Saccade specificity in PPC during the *Visual/Oculomotor Updating* phase. **A.** Upper panels: Voxelwise statistical map from an RFX GLM (n=15) for Different > Same in the *Saccade* condition (orange) is overlaid on inflated cortical surface renderings of right hemisphere of example participant (lateral view on left and medial view on right). Regions in medial precuneus/posterior cingulate gyrus (PCu/pCG) and frontal (MFG, middle frontal gyrus) cortex show saccade-related feature-specific effects. **B.** Specific parietal site localization was performed, where the site-of-interest is shown in white (i.e., ventral PCu, vPCu). The white dot corresponds to location of peak voxel(s) of the site-of-interest. **C.** β-weights were extracted from vPCu, analyzed, and plotted in bar graphs (n=12). In order to meet the first criterion (Prediction 1) to be considered a putative *transsaccadic integrator*, precuneus would need to show a larger feature-related β-weight difference for the *Saccade* and *Fixation* conditions (SD-SS: Saccade Different – Saccade Same; FD-FS: Fixation Different – Fixation Same). As shown in the left bar graph, there is a (significantly) greater feature-related difference for the *Saccade* than for the *Fixation* condition [(SD - SS) > (FD - FS)]. Values are mean differences ± SEM analyzed using one-tailed repeated measures t-tests. (Please see Results for specific statistical values.)

The second step in this analysis was to determine specific parietal coordinates for our hypothesis tests. To do this, we examined the medial PCu/pCG region shown in Fig. 3a, but raised the statistical threshold until only the peak site of parietal activation remained, corresponding to ventral precuneus (vPCu). We then used BrainVoyager (BrainVoyager QX v2.8, Brain Innovation) to determine the cluster of activation around the peak voxel and extract data for prediction testing [24,31,32]. This site is shown in Fig. 3b, overlaid on sagittal anatomic slice (which is less susceptible to distortions). Coordinates and supporting references for this site (and the other regions mentioned) are shown Table 1.

**Table 1.** Names, Talairach coordinates, and sizes of regions of interest (ROIs) observed during the *Visual/Oculomotor Updating* phase.

| Region of Interest | Talairach coordinates |  |  |  |  |  | Active Voxels | References |
| --- | --- | --- | --- | --- | --- | --- | --- | --- |
|  | x | y | z | Std x | Std y | Std z | *n* voxels |  |
| **RH ventral Precuneus** | 14 | -55 | 29 | 1.3 | 1.3 | 2.8 | 142 | [48,91] |
| **RH posterior Cingulate Gyrus** | 9 | -49 | 14 | 2.5 | 2.8 | 2.1 | 641 | [91,92] |
| **RH Middle Frontal Gyrus** | 39 | 17 | 37 | 2.7 | 2.5 | 2.7 | 790 | [93,94] |
| **RH Precentral Gyrus** | 51 | -8 | 34 | 2.8 | 2.7 | 2.8 | 876 | [95] |
| **LH Lingual Gyrus** | -13 | -65 | -11 | 2.4 | 2.4 | 2.6 | 618 | [96] |
| **LH Medial Occipitotemporal Sulcus** | -18 | -77 | -5 | 2.4 | 2.7 | 2.4 | 665 | [97] |

### Hypothesis 1: Saccade-specific feature modulations

According to prediction 1 (Fig. 2), cortical sites involved in transsaccadic feature comparisons should show positive (Different > Same) feature modulations following saccades, but not during sustained fixation. Figure 3a suggests that vPCu is modulated by frequency changes after saccades, but does not show if this modulation is saccade-specific. To test this directly (and independently from the contrast used to localize the site), we extracted β-weights for all participants from a small cubic region surrounding the peak voxel coordinates for vPCu (Table 1). We did this under four conditions for the same site: Saccade Different frequency (SD), Saccade Same frequency (SS), Fixation Different frequency (FD) and Fixation Same frequency (FD). To test hypothesis 1, we then compared *Different* - *Same* feature for *Saccade* (SD-SS) versus *Fixation* data (FD-FS), as shown in Fig. 3c. In short, vPCu followed the predicted pattern: it showed a significantly greater *Different-Same* feature modulation after saccades compared to sustained fixation (t(11)= −2.07, p= 0.031, effect size= 0.60).

### Hypothesis 2: Task-specific saccade modulations

According to prediction 2 (Fig. 2), cortical sites that are specifically involved in transsaccadic feature comparisons should be modulated by saccades in a task-specific fashion, i.e. not simply by the motor act of producing a saccade. Figure 4 provides this test using similar conventions to Figure 3. Figure 4a provides a general overview of the data used in this calculation, showing Saccade > Fixation modulations during the updating portion of our task and for reference, the same contrast from our saccade localizer task, where participants simply looked back and forth between two points (see Methods for details). As in our previous study [24], saccade modulations were generally more widespread when they accompanied a more complex task. Figure 4b shows that the location of the vPCu site used for our hypothesis testing bordered on the edge of a major cluster of saccade modulation.

Figure 4c provides the statistical test of prediction 2, using data derived from our vPCU site in a similar fashion to that described above. i.e. *Saccade-Fixation* difference in the experimental task (TS-TF) versus the same difference from the saccade localizer (S-F). Note that we could only do this for the 12 participants that passed behavioural/imaging criteria in both tasks. Once again these data followed the predicted pattern, showing task-specific saccade modulations in our main experiment compared to the saccade localizer data (t(11)= 2.57, p=0.013, effect size= 0.68). Overall, these findings suggest that activity in our ventral precuneus site reflects changes in spatial frequency in both a saccade-specific and task-specific manner.

**Figure 4.**
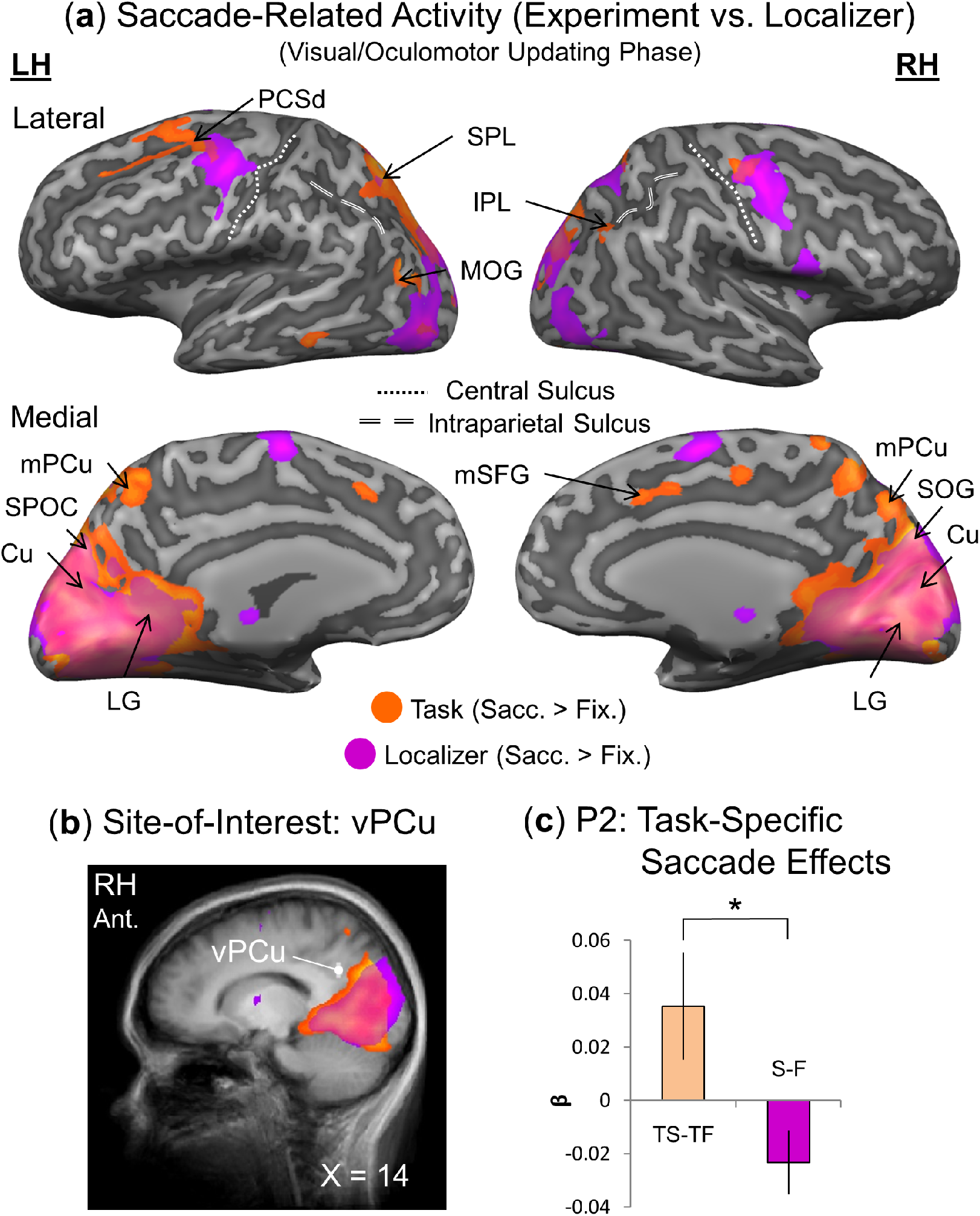
Testing Prediction 2: Task specificity in PPC during the *Visual/Oculomotor Updating* phase. **A.** Upper panels: Voxelwise statistical maps from an RFX GLM (n=15) for Saccade > Fixation in the experimental task (orange; FDR (q < 0.05) and cluster correction) versus the localizer (n=12; fuchsia; FDR (q < 0.05) and cluster correction) are overlaid onto inflated cortical surface renderings of example participant (left hemisphere on the left, right hemisphere on the right; upper panels showing the lateral views, and lower panels showing medial views). There is substantial overlap in activity in medial occipital regions for task and localizer. However, (experimental) task-specific saccade modulations can be observed in particular occipital (middle occipital gyrus, MOG), parietal (medial precuneus, mPCu; inferior parietal lobe, IPL; and superior parietal lobule, SPL), and frontal saccade regions (medial superior frontal gyrus, mSFG – likely pre-supplementary eye field; dorsal precentral sulcus, PCSd – likely frontal eye field). **B.** The site-of-interest (i.e., ventral PCu, vPCu) is shown in white on the sagittal view of the average brain of all the participants, which lies outside of the saccade-specific activation for the task (orange) or localizer (fuchsia). The white dot corresponds to location of peak voxel(s) of the site-of-interest. **C.** β-weights were extracted from vPCu, analyzed, and plotted in bar graphs. To meet the second criterion (Prediction 2), the observed saccade-related modulations for spatial frequency would have to be greater for the task than for a separate saccade-only localizer. The β-weight difference between *Saccade* and *Fixation* conditions is plotted for the task (TS-TF, Task, Saccade and Task, Fixation) and localizer (S-F, Saccade Localizer, Saccade and Saccade Localizer, Fixation). Precuneus also showed a (significant) saccade-related task-specific effect [(TS - TF) > (S – F)]. Values are mean differences ± SEM analyzed using one-tailed repeated measures t-tests. (Please see Results for specific statistical values.)

### Functional network for spatial frequency-specific transsaccadic perception

Our previous study of transsaccadic orientation updating showed a functional network connecting right SMG to both saccade and grasp motor areas [24]. Here, we provide a similar analysis of precuneus activity for the current perceptual task. To do this, we used a *Saccade* > *Fixation* contrast in a psychophysiological interaction (PPI) analysis, wherein right precuneus served as the seed region. Figure 5a shows these data overlaid on a series of transverse brain slices. We observed statistically significant functional connectivity between precuneus (cyan region) and three other regions (orange): 1) left lingual gyrus (LG), 2) a caudal portion of the left medial occipitotemporal sulcus (MOtS), and 3) right precentral gyrus (likely primary motor cortex, M1).

**Figure 5.**
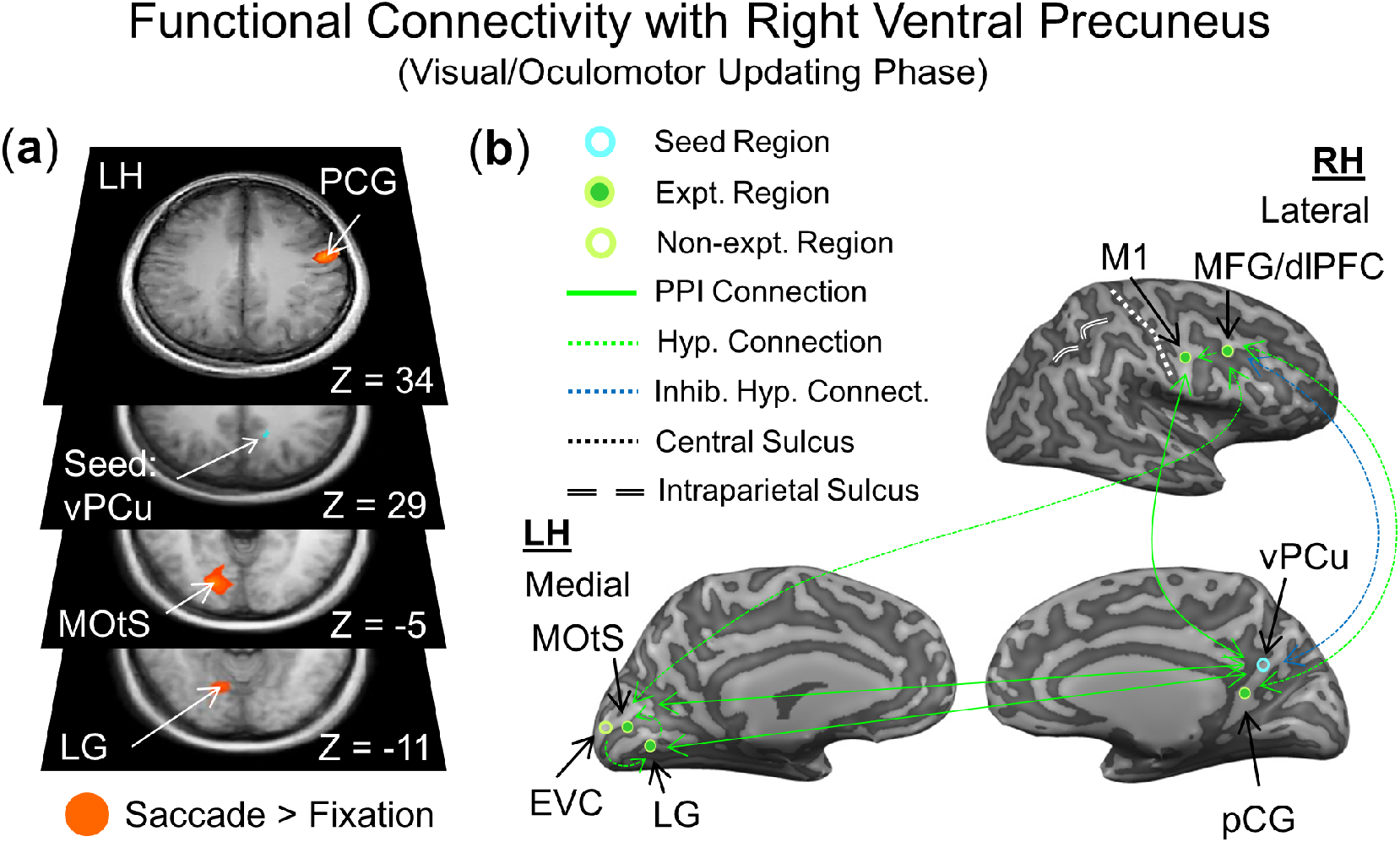
Functional connectivity network (A) and model (B) engaged in transsaccadic updating of spatial frequency. **A.** Transverse slices through the average brain of all participants are shown from most superior to inferior along the vertical axis. ‘Z’ components of each region’s Talairach coordinates are reported for reference. Shown in cyan is the seed region, right precuneus (vPCu). Using a Saccade > Fixation contrast, the psychophysiological interaction (PPI) analysis revealed statistically significant communication between precuneus and right precentral gyrus (PCG, likely primary motor cortex), as well as left lingual gyrus (LG) and medial occipitotemporal sulcus (MOtS). This suggests that parietal precuneus communicates with frontal motor and early/late object perception regions. **B.** A network model proposed to highlight the potential interactions between right precuneus and saccade, memory, and object perception regions. Expt. Region refers to a region that showed activation in our task; Non-expt. Region refers to a region we did not find in our task (e.g., early visual cortex, EVC); PPI connection refers to a connection derived from our PPI results; Hyp. Connection refers to a hypothesized connection between regions in our model; Inhib. Hyp. Connect. refers to the possible inhibitory connection between the medial PCu/pCG region and MFG/dorsolateral prefrontal cortex (dlPFC).

## Discussion

The goal of this study was to determine whether the involvement of PPC in transsaccadic updating of visual location and orientation generalizes to other object features, such as spatial frequency. We found that medial PPC and a lateral frontal region were modulated by spatial frequency changes across saccades. Further hypothesis testing on PPC showed that ventral precuneus passed our predefined criteria for putative transsaccadic updating [24]. Finally, our functional connectivity analysis suggests that this area interacts with other visual and motor areas to perform the task that we required of our participants.

### Putative sites for transsaccadic integration of spatial frequency

It has previously been shown that PPC, specifically intraparietal sulcus and adjacent portions of inferior PPC, are involved in transsaccadic processing of object location and orientation [17,20,24]. The current results extend this role to spatial frequency processing, but implicate a different portion of PPC: ventral precuneus, along with other sites in cingulate and frontal cortex. The latter three sites have several common characteristics. First, although they modulate with saccades, they do not overlap with the classic saccade motor areas (Fig. S2) [33–35]. Second, they were all located in the right hemisphere, consistent with the lateralization observed previously for spatial attention [36,37], spatial updating [38], and transsaccadic orientation processing [20,24]. Finally, they are all relatively ‘high level’ (as opposed to early visual cortex), presumably because our participants performed a top-down, instruction-based task, in contrast to automatic, bottom-up feature integration [11].

Since the intent of this experiment was to explore the role of PPC in transsaccadic integration, we focused our analysis on right ventral precuneus, which met all of our criteria for transsaccadic feature processing. In general, precuneus is associated with complex visuospatial transformations [39–41]. Dorsal precuneus is associated with reaching [42–44], saccades [45,46], and visual memory [47]. Ventral precuneus is involved in change detection [47] and egocentric processing [48], and shows connectivity with the parahippocampus, cuneus, LG (positive connections), and MFG (negative connections) [49]. In short, ventral precuneus has functions and connectivity generally consistent with the task and observations in the current study.

In the current study, precuneus showed extensive functional connectivity with other areas (see next section), and fell within a broad region of right medial posterior cortex activation, that also included another peak in the posterior cingulate gyrus. The posterior cingulate gyrus region is thought to contribute to several saccade-related functions that might be related to transsaccadic perception, including attention orienting [50], and retainment of information in working memory [51]. Saccade-related attention has been closely linked with transsaccadic perception [52].

Finally, we found the opposite pattern of post-saccade activation (*Different* < *Same* frequency) in the right middle frontal gyrus. This is consistent with negative functional connectivity between precuneus and MFG [49]. MFG is located within the dorsolateral prefrontal cortex (dlPFC), which contains various subregions, some saccade-sensitive [53], critical for spatial working memory and top-down control [54–56]. This area has previously been implicated in high-level aspects of transsaccadic perception: e.g., online TMS to dlPFC disrupted memory of multiple objects during fixation, but enhanced transsaccadic memory [57]. In general, dlPFC, has been implicated in various other relevant visual functions, including monochromatic perception of texture [58–60] (although see [61]), attention orientation [62], and retention of perceived information in working memory [55,63,64].

### Putative functional network for transsaccadic perception of spatial frequency

In our previous study of transsaccadic updating of grasp orientation, SMG mainly showed connectivity with sensorimotor areas, including the frontal and supplementary eye fields, anterior intraparietal cortex, and adjacent portions of superior parietal cortex involved in grasp control [24]. Here we observed a very different pattern of connectivity: our seed region (right ventral precuneus) showed task-related functional connectivity with right precentral gyrus (likely M1) [65], left LG, and left MOtS. Although PPI analysis is limited in the sense that it does not provide directionality [66], this analysis suggests a bilateral network for transsaccadic perception of spatial frequency involving both lower and higher-level visual areas. Based on these results, we propose the network model illustrated in Figure 5b, where (as discussed above) ventral precuneus is the central (PPC) hub for spatial frequency processing in this task, and contiguous activity in posterior cingulate contributes saccade-related attention signals [52].

LG (likely corresponding with visual areas V2/V3) processes low-level information about spatial frequency in this network [67–70], and has also been implicated in spatial updating [16,71]. MOtS, located within occipitotemporal cortex, is thought to be involved in higher-level aspects of spatial frequency processing related to objects and scenes [72]. Involvement of precentral gyrus most likely reflects preparatory activity for the final response, the button press [73–75]. Finally, although MFG/dlPFC did not reappear in our connectivity analysis, it was suppressed by feature changes that followed saccades (Fig. 2a) and has been shown to be functionally connected with ventral precuneus in previous studies [49]; a blue dotted line in the model (Fig. 5b) from this area denotes its putative role in top-down control [57,76,77].

### Feature- and task-related aspects of transsaccadic perception

As noted above, our previous experiments showed transsaccadic modulations for object orientation in inferior parietal cortex (specifically SMG) along with some task-dependent extrastriate and superior parietal areas [20,24]. Why would the sites and networks observed for transsaccadic spatial frequency processing in the current study be so different? In part, it is likely that transsaccadic perception builds on computations already used during fixation, which are themselves feature- and task-dependent [21,78,79,80], especially in the higher-level visual areas activated in our tasks [48,65]. At early levels like V1, there is clear multiplexing of orientation and spatial frequency [67,81,82], but at higher levels these may tap into entirely different processes. For example, in our experiment, a change in orientation might influence how one can orient to an object [24,83], but a change in spatial frequency can denote more fundamental changes in perceived identity and affordance, evoking very different cortical mechanisms [84]. Moreover, changes in motor aspects of the task can readily explain why the functional connectivity in the current task (with an abstract button press response) was very different from that in our previous experiment where the perceived object was then grasped [24]. Finally, it is likely that more automatic bottom-up aspects of transsaccadic integration (such as the automatic integration of motion signals across saccades) [85] involve different mechanisms again, perhaps involving earlier visual centres [11,16,71]. Thus, the notion of a dedicated ‘transsaccadic integration centre’ is likely naïve: transsaccadic vision, not prolonged fixation, is normal biological vision, and has likely developed different and nuanced mechanisms, depending on feature and task details.

### Conclusion

In this fMRI study, we set out to explore the cortical mechanism for transsaccadic updating of spatial frequency, with emphasis on the role of PPC. We found that PPC (and likely other cortical areas) play a key role, but in contrast to involvement of SMG for transsaccadic orientation perception [20], the key parietal hub for transsaccadic spatial frequency processing was ventral precuneus. We also found different patterns of functional connectivity compared to our previous grasp updating experiment, i.e., where SMG activity correlated with saccade and grasp motor areas [24]. Overall, these findings support the role of parietal cortex in transsaccadic vision, but suggest that specific network recruitment is feature- and task-dependent.

## Materials and Methods

### Participants

We tested 21 (human) participants from York University (Toronto, Canada) of whom 15 met our inclusion criteria for analyses (see Behavioural analysis and exclusion criteria), thus exceeding the requirement for sufficient statistical power (see Power analysis) [86]. These 15 individuals (average age: 26.6 +/- 4.3 years; age range: 21-37; 11 females and 4 males; all right-handed) had no neurological disorders and normal or corrected-to-normal vision. Informed consent was obtained from each participant; all participants were remunerated for their time. We confirm that the experiment protocol involving human participants was approved by and in accordance with guidelines of the York University Human Participants Review Subcommittee.

### Experimental set-up

Participants passed initial MRI safety screening and were then informed about the experimental task. Participants practiced before taking part in the experiment. Once they felt comfortable with the task, they assumed a supine position on the MRI table, with their head resting flat within a 32-channel head coil. An apparatus, holding a mirror to reflect images from the screen in the MRI bore, was attached to the head coil. An MRI-compatible eye-tracker (iViewX, SensoMotoric Instruments) was also attached to the apparatus in order to record the position of the right eye. Participants held an MRI-compatible button box in their right hand. Their right index and middle fingers rested on two buttons, used to provide the task responses when presented with the ‘go’ cue (see General paradigm).

### Stimuli

During the experiment, participants were presented with an 18° stimulus that contained a vertical sine-wave grating pattern, averaged to the mean luminance of the screen. Stimuli were presented in the centre of the screen on a light gray background (MATLAB; Mathworks, Inc.). The two spatial frequencies that were tested were 0.7 or 1.1 cycles/degree (cpd) (see Fig. 1a).

### General paradigm

In order to identify the cortical activity correlates of transsaccadic perception of spatial frequency, we used a modified 2 (Gaze: Fixate or Saccade) x 2 (Spatial Frequency: 0.7 or 1.1 cpd) slow event-related fMRI design [20]. Here, we modulated the spatial frequency (repeated or changed within the same trial; *‘Same’* and *‘Different’* conditions, respectively) and/or position of the eyes (continued fixation of gaze or a saccade was produced; *‘Fixation’* and *‘Saccade’* conditions). This resulted in four main conditions: 1) Fixation/Same, 2) Fixation/Different, 3) Saccade/Same, and 4) Saccade/Different. These were randomly intermingled and repeated four times within each run; there were six runs in total for each participant.

### Trial sequence

Each trial was composed of three main phases: 1) an initial presentation of the stimulus, which requires sensory processing and working memory storage (‘ *Sensory/Memory’* phase); 2) a second presentation of the stimulus, which requires sensory processing and (oculomotor) updating (for saccades) (*‘Visual/Oculomotor Updating’* phase); and 3) a response period, where a button press was made to indicate if the spatial frequency of the stimulus presentations was the same or different (*‘Response’* phase). The sequence of events in each trial (Fig. 1a) began with a 2 s fixation period, with the fixation cross presented at one of two possible positions (13.5° to the left or right of centre, or 4.5° to the left or right of the central stimulus). The stimulus was then presented in the centre for 5.8 s, followed by a 200 ms static noise mask. Once the mask disappeared, a fixation cross appeared for 200 ms at the same position (*Fixation* condition) or the other fixation position (*Saccade* condition) (see Fig. 1b for eye position trace). The central stimulus was presented a second time for 5.8 s at the same spatial frequency (*Same* condition) as in the *Sensory/Memory* phase or at the other spatial frequency (*Different* condition). Finally, after the fixation cross and stimulus disappeared, a written prompt (‘R or N?’) appeared for 8 s in the same location as the fixation cross, prompting participants to indicate via button press whether the spatial frequency of the two stimulus presentations was the same (R) or different (N).

Each trial sequence lasted 22 s in total. The four main conditions were repeated four times per run, resulting in 16 trials per run. Each run started and ended with a 16 s period of central fixation, serving as a baseline. Overall, each run lasted 6 min 24 s.

### Saccade localizer

We ran a separate saccade localizer in order to identify regions of the brain that respond to saccade production. The sequence of events began with an 8 s fixation period of a central white cross on a black background. This was followed by a period of directed saccadic eye movements between two fixation crosses, randomly from left to right across trials, for 8 s. The pattern of fixation followed by saccades was repeated 16 times per localizer run, lasting ~4 min 16 s for each of two runs.

### MRI parameters

A 3T Siemens Magnetom TIM Trio MRI scanner at the York MRI Facility was used to acquire fMRI data. An echo-planar imaging (EPI) sequence (repetition time [TR] = 2000 ms; echo time [TE] = 30 ms; flip angle [FA] = 90°; field of view [FOV] = 240 x 240 mm, matrix size = 80 x 80, in-plane resolution = 3 mm × 3 mm; slice thickness = 3 mm) was acquired in ascending, interleaved order for each of the six functional runs and for the two separate saccade localizer runs. Thirty-three contiguous slices were acquired per volume for a total of 192 volumes of functional data in each experimental run, and 128 volumes of data for each saccade localizer run. A T1-weighted anatomical reference volume was obtained for each participant using an MPRAGE sequence (TR= 1900 ms; FA= 256 mm x 256 mm; voxel size= 1 x 1 x 1 mm^3^). 192 slices were acquired per volume of anatomical data.

### Analysis

#### Power analysis

To determine the appropriate number of participants to provide a sufficient effect size and level of power, we used results from the most recent, relevant findings [24]. Specifically, we used the effect size (0.887, Cohen’s d) from the most relevant region of activation in parietal cortex (i.e., SMG) and applied the following properties: 1) two-tailed t-test option, 2) an α value of 0.05, and 3) a power value of 0.85. Using G*Power 3.1 [87], we determined that a minimum of 14 participants would be required to achieve an actual power value of 0.866. As noted above, we tested a total of 21 participants in order to exceed this minimal number, after application of exclusion criteria in the next section.

#### Behavioural data and exclusion criteria

In our previous studies [20,24], we found that experiments such as this are highly sensitive to head motion. Due to excessive head motion, data were excluded from analyses on the basis of two criteria: 1) presence of head motion over 1 mm/degree and/or 2) any abrupt motion of the head over 0.5 mm/degrees. If more than 50% of all data (i.e., at least 3 runs out of the total 6 runs for a given participant) were removed from analysis, the entire data set from that participant was removed. On this basis, data from six participants were removed. From the remaining 15 participants, one run was removed from data analysis for each of two participants, and two runs were removed for each of another two participants, for a total of six runs (6.7%).

Eye tracking and button-press data were analyzed post-image acquisition in order to determine whether the task was completed correctly or not. Eye position data (e.g., Fig. 1b) were inspected visually to confirm that the eye fixated on the fixation crosses and/or moved to the correct saccade location when prompted to do so in all trials. Button press responses were also inspected offline to ensure that participants responded correctly to the *Same/Different* condition trials. On these bases, 41 trials were excluded from further analysis (3.1% of the remaining data). Overall, accuracy of the 15 participants included in the final data analysis was 97.4% ± 3.1%. Only data for correct trials were included in all further analyses.

#### Functional imaging data: experimental

To model the fMRI BOLD response, we used a general linear model (GLM) analysis. In this model, a standard two-gamma haemodynamic response function (BrainVoyager QX 2.8, Brain Innovation) was convolved with predictor variables [20]. We had five major classes of predictors for each trial: 1) a baseline predictor (“Baseline”), corresponding to the first and last 16 s of each run; 2) “Fixate”, which represented initial trial fixation (either left or right); 3) “Adapt”, which modeled the activity in response to the first stimulus presentation; 4) “Fixate/Same”, “Fixate/Different”, “Saccade/Same”, or “Saccade/Different” to model activity in response to the second stimulus presentation in one of the four main conditions; and 5) “Response”, which modeled the activity for the button press response period. GLMs were generated using the eight predictors per run for each participant (BrainVoyager QX 2.8, Brain Innovation).

Preprocessing of functional data from each run for all participants included slice scan-time correction (cubic spline), temporal filtering (for removal of frequencies < 2 cycles/run), and 3D motion correction (trilinear/sinc). Anatomical data were transformed to a standard Talairach template [88]. Functional data were coregistered using gradient-based affine alignment (translation, rotation, scale affine transformation) to the raw anatomical data. Lastly, the functional data were spatially smoothed using a Gaussian kernel with a FWHM of 8 mm.

Using the random-effects (RFX) GLM with all of the runs of all of the remaining 15 participants, we performed two major types of analyses: 1) parietal site-of-interest prediction testing and 2) voxelwise contrasts (see Hypothesis testing: Analysis and statistical considerations and Voxelwise map contrasts: Analysis and statistical considerations for details).

#### Functional imaging data: saccade localizer

Each run of the saccade localizer had 16 repetitions of the fixation trials (8 s of central fixation) and saccade trials (8 s of directed saccades). Each trial type was coded by an 8 s predictor (“Fixation” for the fixation trials, and “Saccade” for the saccade trials). These predictors were convolved with the standard two-gamma haemodynamic response function (BrainVoyager QX 2.8, Brain Innovation). Preprocessing of functional and anatomical data occurred as for experimental data (see previous).

On the basis of behavioural data analysis (i.e., excessive motion > 1 mm), data from three participants were removed, leaving saccade localizer data for 12 participants. These data were used in the prediction testing in order to identify regions related to the production of saccades.

#### Voxelwise map contrasts: Analysis and statistical considerations

We conducted voxelwise contrasts on data from the 15 participants included in the analysis (for task, and on the remaining 12 participants for the localizer). For analysis purposes, we divided the experimental task data into the *Sensory/Memory* phase (when participants saw and initially remembered the stimulus) and *Visual/Oculomotor Updating* phase (when saccades occurred and the stimulus reappeared). For all (voxelwise) contrasts, we first applied a False Discovery Rate (FDR) of q < 0.05, followed by cluster threshold correction (BrainVoyager QX v2.8, Brain Innovation) to our contrasts. This included the Saccade > Fixation contrast shown in Fig. 4a/b, and the contrasts shown in supplementary figures (Sensory/Memory phase activity > Baseline, Fig. S1; Visual/Oculomotor Updating > Sensory/Memory contrast, Fig. S2).

#### Hypothesis testing: Analysis and statistical considerations

In order to localize our site-of-interest testing, we used a similar approach to site localization as in Song and Jiang [31]; Baltaretu et al. [24]; and Tsushima et al. [32] by first applying a whole-brain contrast of interest: Saccade Different – Saccade Same at a liberal p-value (p < 0.05) with cluster threshold correction (Fig. 3a). We then used reduced the p-value in order to localize the centre of our *a priori* predicted parietal activation (i.e., within the larger pCU/pCG area, Fig. 3b), and then used BrainVoyager (BrainVoyager QX v2.8, Brain Innovation) to determine a cluster within a 1000 mm^3^ volume (p < 0.05, cluster corrected; Table 1). Finally, we extracted the β-weights from associated voxels around the peak voxel for the experimental data and saccade localizer data (separately) in order to test our hypotheses within the parietal, ventral precuneus (i.e., first, to determine the presence of a transsaccadic feature-specific effect and, second, whether transsaccadic effects are task-specific). These analyses were carried out for the 12 participants who survived our behavioural criteria for the task and localizer.

Using the β-weights, we looked for specific directionality in the two predictions that we tested (Fig. 2), so we used one-tailed repeated measures t-tests for each of the two predictions (one to identify saccade/feature-related specificity and another to determine transsaccadic task specificity). For the results of these analyses, we provided all relevant t-statistics, p-values, and effect sizes (Cohen’s d, determined using G*Power) [87].

#### Psychophysiological interaction

Lastly, we were interested in uncovering the functional network for saccade-related updating during transsaccadic perception. In order to do this, we conducted psychophysiological interaction (PPI) analysis [24,66,89,90] on data extracted from the posterior parietal region that showed transsaccadic effects from the *Visual/Oculomotor Updating* phase. To perform this analysis, we used three predictors: 1) a physiological component (which is accounted for by z-normalized time courses obtained from the seed region for each participant for each run); 2) a psychological component (task predictors, i.e. for the Saccade and Fixation conditions, were convolved with a hemodynamic response function), and 3) a psychophysiological interaction component (multiplication of seed region z-normalized time courses with task model in a volume-by-volume manner). For the psychological component, the *Saccade* predictors were set to a value of ‘+1’, whereas the Fixation predictors were set to a value of ‘-1’; all other predictors were set to a value of ‘0’ in order to determine the responses to Saccades as compared to Fixation. (For greater detail, please see O’Reilly et al. [66] and Baltaretu et al. [24].) We created single design matrices for each run for each of the 15 participants, which were ultimately included in an RFX GLM. Then, we used the third, psychophysiological interaction predictor to determine the regions that comprise the functional network that show saccade-related modulations for feature updating. We applied the p-value threshold (0.05) to the volumetric map and cluster threshold correction to identify clusters of activation. We used similar region localization as Song and Jiang [31], Baltaretu et al. [24], and Tsushima et al. [32] to pinpoint specific regions of activation for this analysis.

## Acknowledgements

The authors thank Joy Williams, Dr. Xiaogang Yan, and Saihong Sun for technical assistance. This work was supported by a Natural Sciences and Engineering Council (NSERC) of Canada Discovery grant. B.R. Baltaretu was supported by the NSERC Brain-in-Action CREATE program, and J.D. Crawford is supported by the Canada Research Chair Program.

## Author Contributions

B.R.B. adapted the paradigm, collected and analyzed the data, and wrote the manuscript. B.T.D. developed the original paradigm and edited the manuscript. W.D.S. supervised data analysis and edited the manuscript. J.D.C. supervised overall project development and edited the manuscript.

