## Supplementary Figs and Info for "Parietal Mechanisms for Transsaccadic Spatial Frequency Perception: An fMRI Study" (from another version of this preprint: version 1, 2020-07-15)

### Supplementary Materials and Methods

#### Regions of interest.

**Contrasts to determine regions active during the Sensory/Memory Phase.**

In order to understand how the brain processes spatial frequency information in preparation of transsaccadic perception, we applied a contrast (Task > Baseline; FDR ($q < 0.05$) to the volumetric map produced for data from the *Sensory/Memory* phase (Fig. 1a). During this phase, we anticipated activity in regions in 1) occipital cortex that are involved in early visual processing of the spatial frequency stimulus, 2) parietal regions for the spatial encoding of feature information, and 3) frontal, saccade-related regions involved in producing visually-guided eye movements. Regions that were statistically significant and survived cluster threshold correction are labeled and further discussed (see Results and Discussion).

**Contrasts to determine change in activation from Sensory/Memory to Visual/Oculomotor Updating phase.**

We applied a voxelwise contrast (Sensory/Motor > Visual/Oculomotor Updating phase; FDR ($q < 0.05$)) in order to identify the regions of activation associated with each of the two stimulus presentation phases of our task. During the *Sensory/Memory* phase, we expected to observe activation in 1) early occipital regions, as well as some 2) object-memory temporal regions. During the *Visual/Oculomotor Updating* phase, we predicted that we would observe activation in 1) later occipital, 2) sensorimotor parietal, and 3) saccade-related frontal regions. Regions that were statistically significant and survived cluster threshold correction are labeled and further discussed (see Results).

### Supplementary Results

#### General task-related activation (Sensory/Memory phase).

Here, we first used a contrast (*Sensory/Memory* Activity > Baseline contrast) during the *Sensory/Memory* phase (i.e., when the stimulus was first presented) to determine cortical activation involved in the initial processing spatial frequency of the stimulus. Given the mechanisms required during this phase in our task (i.e., sensory processing of the visual stimulus, spatial encoding for potential oculomotor updating in the case of saccades, and retainment of characteristic of the stimulus for later comparison), we anticipated that regions within 1) occipital (sensory processing), 2) parietal (spatial encoding), and 3) frontal (saccade production) cortex would be active [10,23,35,66].

The cortical regions that were activated in response to the first presentation of the spatial frequency stimulus during the *Sensory/Memory* phase covers the hypothesized areas of the cortical cortex (Fig. S1). When we specifically localized the regions within the larger clusters of activation across the cortex, within occipital cortex, we found right lingual gyrus (LG), left inferior occipital gyrus (IOG), and bilateral middle occipital gyrus (MOG). In parietal cortex, we found activation with the left superior parietal lobule (SPL), at the junction with the parieto-occipital sulcus (POS). Lastly, there was also frontal lobe activation within left dorsal precentral sulcus (PCSd; likely frontal eye field, FEF) and superior frontal gyrus (SFG; likely supplementary eye field, SEF). Thus, when a spatial frequency stimulus is observed initially, especially when it precedes a potential updating (change in feature and/or change in gaze), an extensive swath of regions that are involved in early sensory processing, spatial encoding, and eye-related modulations are engaged.

**Phase-related cortical activation differences.**

In order to determine how cortical activation changes across the two stimulus presentation phases, we used a voxelwise contrast (i.e., Sensory/Memory > Visual/Oculomotor Updating activity) to do this. In light peach on the inflated brain rendering of an example participant (Fig. S2) are regions that show greater activation during the first, *Sensory/Memory* phase as compared to the *Visual/Oculomotor Updating* phase. These regions are found in the medial parieto-limbic PCu/pCG region, as well as in more anterior superior frontal gyrus (aSFG), which are not the early occipital/temporal regions we would have expected to see here. On the other hand, regions in burnt orange that showed greater sensitivity to the *Visual/Oculomotor Updating* phase as compared with the *Sensory/Memory* phase were observed in early visual/occipital, parietal cortex (pIPS and SPL), as well as in saccade-related frontal regions. These were in line with our expectations for this direction of the phase contrast. This suggests that there is a transition or preference for different parietal/frontal regions during the first stimulus presentation period, compared to the vast activation extending across occipital, parietal, and frontal-saccade cortex.

### Supplementary Figures

### Figure Legends

**Figure S1.** General task-related activity derived from the *Sensory/Memory* phase. On the inflated brain renderings of an example participant (lateral views shown in the top panels, medial views shown in the bottom panels) in the left and right hemispheres (shown on the left and right, LH, RH, respectively) is overlaid activity (n=15) for the first stimulus presentation (contrast: Task > Baseline). As anticipated, early visual (inferior occipital gyrus, middle occipital gyrus, and lingual gyrus), spatial-encoding parietal (superior parietal lobule), and eye-movement frontal (dorsal precentral sulcus (likely frontal eye field) and superior frontal gyrus (likely pre-/supplementary eye field)) showed activation in response to the initial stimulus presentation. *Abbreviations*: IOG, inferior occipital gyrus; MOG, middle occipital gyrus; LG, lingual gyrus; SPL, superior parietal lobe; PCSd, dorsal precentral sulcus; mSFG, medial superior frontal gyrus.

**Figure S2.** Phase-specific cortical activation differences for the *Sensory/Memory* and *Visual/Oculomotor Updating* phases. On the inflated brain rendering of an example participant (left panels represent left hemisphere, right panels represent right hemisphere; upper panels show lateral views, lower panels show medial views) are the voxelwise statistical maps of the Sensory/Memory (S/M) > Visual/Oculomotor Updating (V/O) contrast. Regions that show greater sensitivity during the *Sensory/Memory* phase over the *Visual/Oculomotor Updating* phase are shown in light peach; this activation is observed in medial parieto-limbic cortex (PCu/pCG) and in anterior frontal regions (aSFG). In contrast, regions in burnt orange show greater sensitivity for the change occurring in the *Visual/Oculomotor Updating* phase than in the *Sensory/Motor* phase. This activation spans occipital, parietal, and saccade-related frontal cortex. This suggests a difference in cortical activation across the two stimulus presentation phases. *Abbreviations*: LG, lingual gyrus; Cu: cuneus; IOG, inferior occipital gyrus; PCu/pCG, precuneus/posterior cingulate gyrus; pIPS, posterior intraparietal sulcus; SPL, superior parietal lobule; PCG, precentral gyrus; PCSd, dorsal precentral sulcus; mSFG, medial superior frontal gyrus; aSFG, anterior superior frontal gyrus.

**Figure S1**

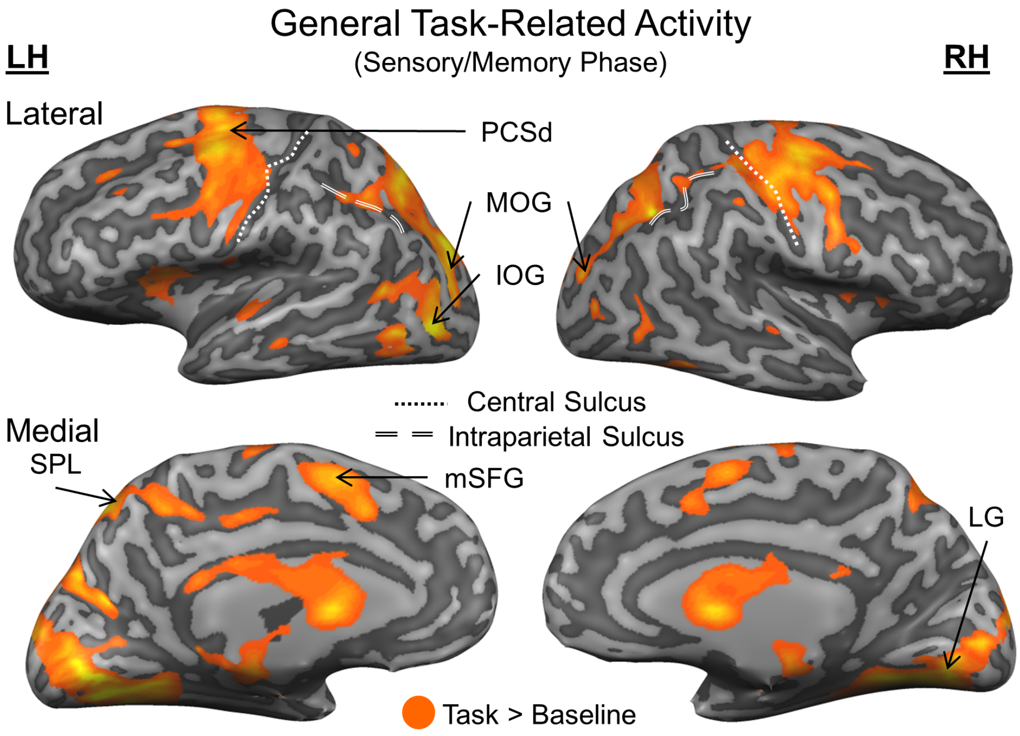
General Task-Related Activity
(Sensory/Memory Phase)
LH
RH
Lateral
PCSd
MOG
IOG
Central Sulcus
Intraparietal Sulcus
Medial
SPL
mSFG
LG
Task > Baseline

**Figure S2**

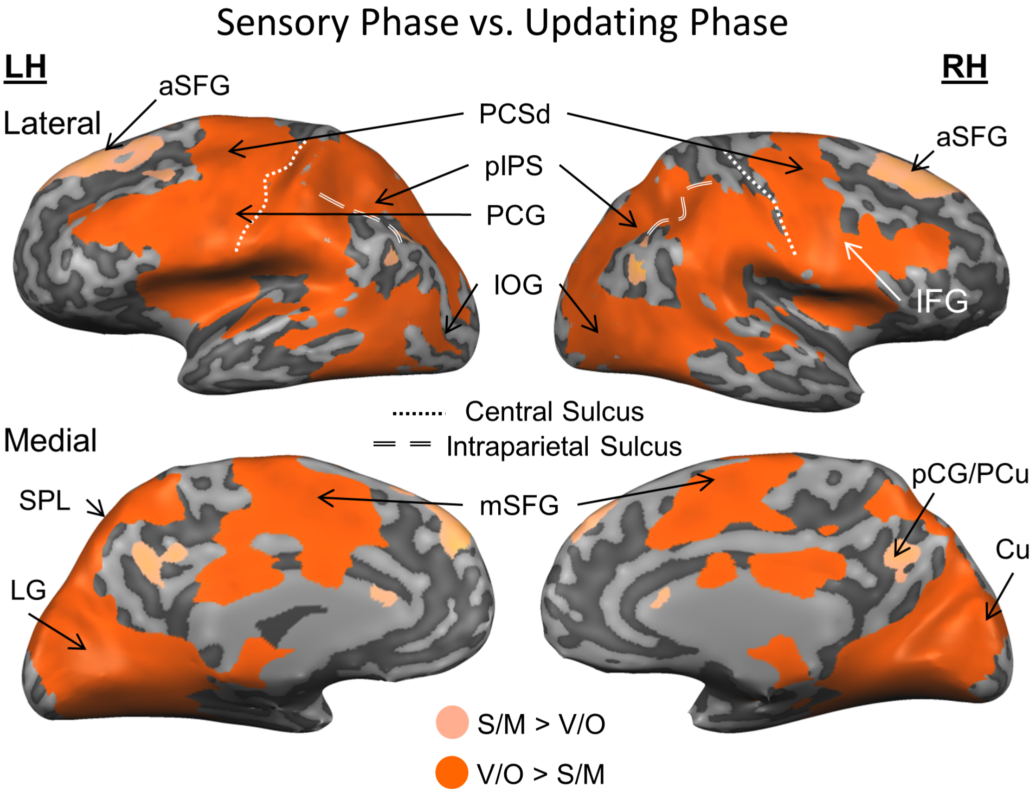
Sensory Phase vs. Updating Phase
LH
RH
Lateral
aSFG
PCSd
pIPS
PCG
IOG
aSFG
IFG
Central Sulcus
Intraparietal Sulcus
Medial
SPL
LG
mSFG
pCG/PCu
Cu
S/M > V/O
V/O > S/M
